# An epigenome-wide association study of sex-specific chronological ageing

**DOI:** 10.1101/606020

**Authors:** Daniel L. McCartney, Robert F. Hillary, Qian Zhang, Anna J. Stevenson, Rosie M. Walker, Mairead L. Bermingham, Stewart W. Morris, Archie Campbell, Alison D. Murray, Heather C. Whalley, David J. Porteous, Kathryn L. Evans, Tamir Chandra, Ian J. Deary, Andrew M. McIntosh, Peter M. Visscher, Allan F. McRae, Riccardo E. Marioni

## Abstract

**Introduction:** Advanced age is associated with cognitive and physical decline, and is a major risk factor for a multitude of disorders including neurodegenerative diseases such as Alzheimer’s disease. There is also a gap in life-expectancy between males and females. DNA methylation differences have been shown to be associated with both age and sex. Here, we investigate age-by-sex differences in DNA methylation in an unrelated cohort of 2,586 individuals between the ages of 18 and 87 years.

**Methods:** Genome-wide DNA methylation was measured on the Illumina HumanMethylationEPIC beadchip in a subset of unrelated individuals from the Generation Scotland cohort. Mixed linear model-based analyses were performed to investigate the relationship between DNA methylation and an interaction term between age and sex, as well as chronological age.

**Results:** At a genome-wide significance level of *P* < 3.6 × 10^−8^, 14 loci were associated with the age-by-sex interaction term, the majority of which were X-linked (n = 12). Seven of these loci were annotated to genes. The site with the greatest difference mapped to *GAGE10*, an X-linked gene. Here, DNA methylation levels remained stable across the male adult age range (DNA methylation x age r = 0.02), but decreased across female adult age range (DNA methylation x age r = −0.61). The seven age-by-sex-associated genes were enriched among differentially-expressed genes in lung, liver, testis and blood.

**Conclusion:** The majority of differences in age-associated DNA methylation trajectories between sexes are present on the X-chromosome. Several of these differences occur within genes which have implicated in multiple cancers, schizophrenia and systemic lupus erythematosus.

## Introduction

Advanced age is associated with cognitive and physical decline, and is a major risk factor for a multitude of disorders including cancer, cardiovascular disease and neurodegenerative diseases. Biological hallmarks of ageing have been observed at the cellular and molecular level and include shortening of telomeres, genomic instability and both global and local changes in DNA methylation (DNAm) levels [1, 2, 3]. DNAm is a common epigenetic mark, typically occurring in the context of a Cytosine-Guanine dinucleotide motif (CpG). It can be modulated by both environmental exposures and genetic variation, and recent work has developed robust predictors of chronological age-based DNAm profiles at subsets of loci (the epigenetic clock) [4, 5].

The average life expectancy at birth in Scotland is 77.0 years for males and 81.1 years for females as reported by the Life Tables for Scotland 2015-2017 [6]. This disparity may be explained in part by differences in biological ageing mechanisms between males and females, and is supported by reports of an increased DNAm-based age acceleration in males relative to females (i.e. a ‘faster ticking’ epigenetic clock) [7]. In the current study, we sought to identify sex differences in age-associated genome-wide DNAm changes, in a cohort of 2,586 unrelated individuals. Further understanding of the sex-specific effects on biological ageing may assist in identifying novel risk factors for age- and sex-associated pathologies.

## Methods

### Generation Scotland: Scottish Family Health Study

Data came from the family-based Generation Scotland: Scottish Family Health Study (GS). GS participants were recruited from GP practices in five regions across Scotland between the years 2006 and 2011 [8]. The probands were aged between 35 and 65 years and were asked to invite first degree relatives to join the study, which had a final size of 24,090. A variety of cognitive, physical, and health data were collected at the study baseline along with blood or saliva samples for DNA genotyping. Blood-based DNAm data were obtained on a subset of 5,200 participants using the Illumina EPIC array [9]. Quality control details have been reported previously [9]. Briefly, probes were removed based on: (i) outliers from visual inspection of the log median intensity of the methylated versus unmethylated signal per array; (ii) a beadcount <3 in more than 5% of samples and; (iii) ≥0.5% of samples having a detection p-value >0.05. Samples were removed (i) if there was a mismatch between their predicted sex and recorded sex and/or (ii) if ≥1% of CpGs had a detection p-value >0.05. For the present analyses, we considered unrelated individuals from the DNAm subset of GS. A genetic relationship matrix was built using GCTA-GRM and a relatedness coefficient of <0.025 was specified to exclude related individuals [10]. In cases where a couple was present, one individual was removed to minimise shared-environment effects. This left an analysis sample of 2,586 unrelated individuals ranging in age from 18 to 87 years and 807,857 probes.

### Ethics

All components of GS received ethical approval from the NHS Tayside Committee on Medical Research Ethics (REC Reference Number: 05/S1401/89). GS has also been granted Research Tissue Bank status by the East of Scotland Research Ethics Service (REC Reference Number: 15/0040/ES), providing generic ethical approval for a wide range of uses within medical research.

### Statistical analysis

All analyses were performed in R version 3.4.2 [11].

### Epigenome-Wide Association Studies

Epigenome-wide association studies (EWASs) of chronological age and the interaction between age and sex were performed using OmicS-data-based Complex trait Analysis software (OSCA), adjusting for smoking status and estimated cell proportions (CD8+ T-cells, CD4+ T-cells, natural killer cells, B-Cells, and granulocytes [5]). DNAm was the outcome variable, age, and the interaction between age and sex, were the predictor variables of interest. Age was scaled to have mean 0 and variance 1. The MOMENT method was used to test for associations between the traits of interest and DNAm at individual probes [12]. MOMENT is a mixed-linear-model-based method that can account for unobserved confounders and the correlation between distal probes which may be introduced by such confounders. Probes with a P-value less than 3.6×10^−8^ [13] were considered epigenome-wide significant associations.

### Gene expression analysis

Gene expression data from 369 whole-blood samples were downloaded from the Genotype-Tissue Expression (GTEx) project (https://gtexportal.org/home/) [17]. Age information was present as six 10-year bins (20-29, 30-39, 40-49, 50-59, 60-69, and 70-79). Samples were stratified by sex and Spearman’s correlations between median age per bin and expression were assessed for genes identified as genome-wide significant in the age-by-sex interaction EWAS. To permit comparison with the DNAm dataset, Ensembl gene IDs were converted to gene symbols using the *org.Hs.eg.db* package [14].

### Pathway analysis

Gene symbols as annotated by the *IlluminaHumanMethylationEPICanno.ilm10b2.hg19* package were converted to Entrez IDs using the *org.Hs.eg.db* package [14]. Of the 26,737 annotated genes, 23,630 were successfully converted to Entrez IDs (88.4%). Enrichment was assessed among KEGG pathways and Gene Ontology (GO) terms via hypergeometric tests using the *phyper* function in R. KEGG pathway data were acquired from the KEGG REST server on October 01 2018. GO data were acquired from the C5 gene set collection from MSigDB [15, 16].

## Results

### Sample demographics

The genetically unrelated cohort subset had a mean age of 50 years (SD = 12.5) and comprised 1,587 females (61.4%) and 999 males (38.6%). Males ranged in age from 18.1 years to 85.7 years (mean = 50.8 years, SD = 12.2) whereas females ranged from 18.0 years to 86.9 years (mean = 49.5 years, SD = 12.7).

### DNAm and chronological age

There were 989 CpGs associated with chronological age at the epigenome-wide significant level (*P* < 3.6×10^−8^; **Supplementary Figure 1**), annotated to 557 unique genes. The top associations mapped to sites previously associated with chronological age in the genes *KLF14, ELOVL2* and *FHL2* (*P* < 1×10^−112^; univariate Spearman correlations with age ranged between 0.59 and 0.75) [4, 5, 18]. A summary of all epigenome-wide significant CpG associations with chronological age is available in **Supplementary Table 1.**

### DNAm and the interaction between chronological age and sex

There were 14 epigenome-wide significant CpG associations with the age-sex interaction term, annotated to seven unique gene symbols (*GAGE10, ANKRD58, CXorf38, TMEM164, RENBP, MID2*, and *OTUD5*), all of which were X-linked. Twelve of the 14 CpGs (85.7%) were on the X-chromosome, one of which (cg15833111 [*GAGE10*]) was also significantly associated with chronological age (*P* = 3.8 × 10^−10^; **Figure 1; Table 1**).

**Figure 1:**
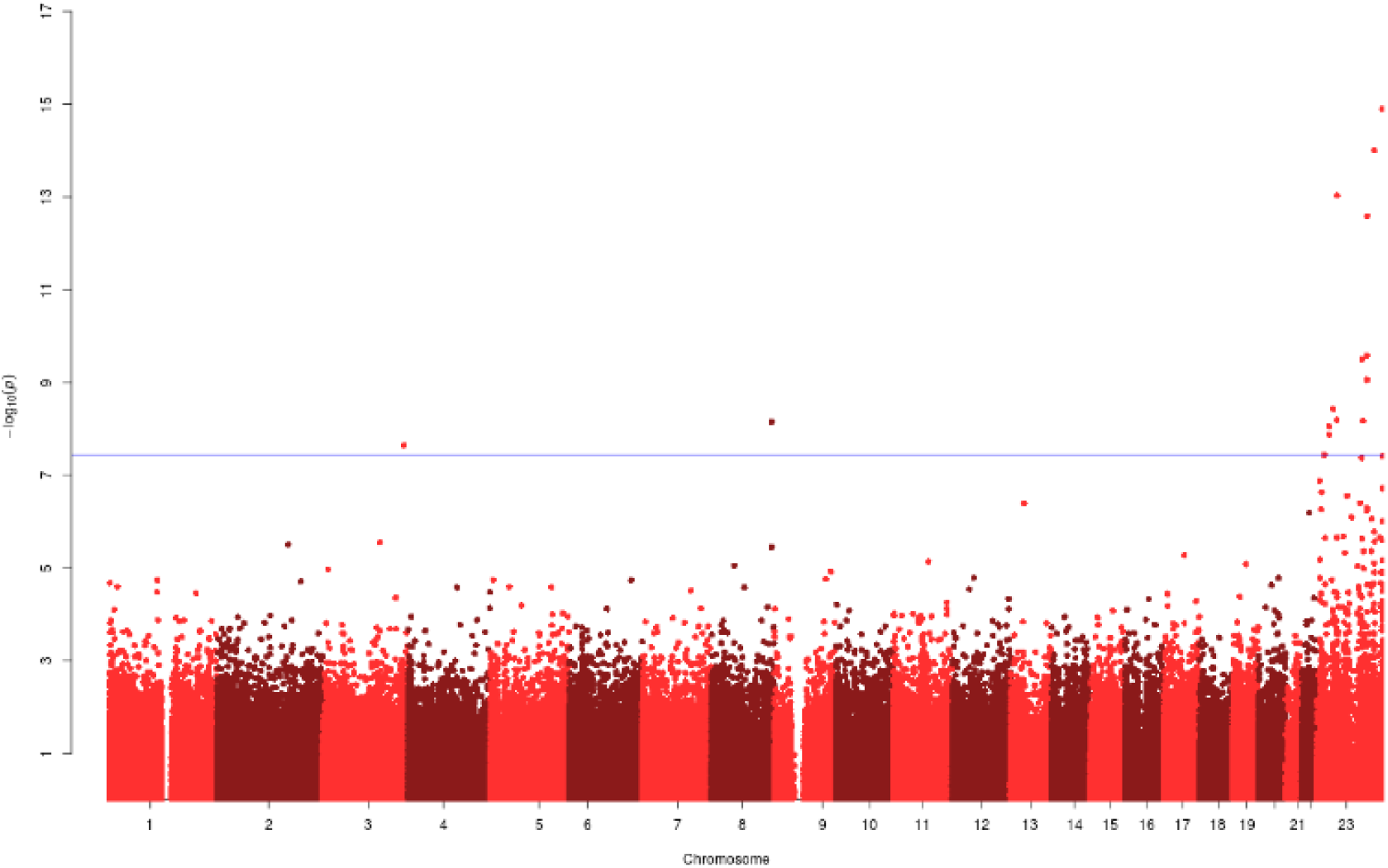
Manhattan Plot for the EWAS of the age-by-sex interaction in Generation Scotland.

**Table 1.** Genome-wide significant (P<3.6×10^−8^) age x sex interaction CpG sites

| Probe | Chr | Bp | Gene | Feature | Beta | SE | P | Chronological Age?* |
| --- | --- | --- | --- | --- | --- | --- | --- | --- |
| cg16464950 | X | 1.53E+08 | *RENBP* | ExonBnd;Body | 0.21 | 0.03 | $1.26 \times 10^{-15}$ | N |
| cg24541420 | X | 1.36E+08 | | | 0.13 | 0.02 | $9.97 \times 10^{-15}$ | N |
| cg15833111 | X | 49166019 | *GAGE10* | Body | -0.14 | 0.02 | $9.28 \times 10^{-14}$ | Y |
| cg13217333 | X | 1.19E+08 | *ANKRD58* | TSS1500 | 0.14 | 0.02 | $2.59 \times 10^{-13}$ | N |
| cg20202246 | X | 1.18E+08 | | | -0.16 | 0.03 | $2.65 \times 10^{-10}$ | N |
| cg04828704 | X | 1.07E+08 | *MID2* | Body | 0.13 | 0.02 | $3.15 \times 10^{-10}$ | N |
| cg08814148 | X | 1.18E+08 | | | -0.10 | 0.02 | $8.80 \times 10^{-10}$ | N |
| cg06231618 | X | 40504059 | *CXorf38* | Body | 0.15 | 0.02 | $3.77 \times 10^{-09}$ | N |
| cg03711046 | X | 48814205 | *OTUD5* | Body;5'UTR | 0.12 | 0.02 | $6.44 \times 10^{-09}$ | N |
| cg07911663 | X | 1.09E+08 | *TMEM164* | TSS1500 | 0.15 | 0.03 | $6.75 \times 10^{-09}$ | N |
| cg11530334 | 8 | 1.4E+08 | | | 0.06 | 0.01 | $7.05 \times 10^{-09}$ | N |
| cg05548968 | X | 30928416 | | | -0.09 | 0.02 | $8.87 \times 10^{-09}$ | N |
| cg25140188 | X | 31087348 | | | 0.08 | 0.01 | $1.32 \times 10^{-08}$ | N |
| cg06123699 | 3 | 1.89E+08 | | | -0.18 | 0.03 | $2.24 \times 10^{-08}$ | N |

\* This column indicates whether or not DNA methylation at the corresponding probe is also associated with chronological age.

The probe with the greatest ageing-associated difference between males and females was cg15833111, mapping to *GAGE10* on the X chromosome (Figure 2; interaction P = 9.3 × 10^−14^). Hypomethylation of this probe was observed with increasing age in females (r = −0.61), whereas methylation levels remained stable in males (r = 0.02). Nine of the 14 CpG sites exhibited clear differences in mean DNAm levels between males and females across an adult age range. These ranged from 0.14 to 0.34 on the beta-value scale, with females displaying consistently higher methylation levels. The remaining probes showed progressive hypomethylation across the adult age range in females (**Figure 2**); hyper-, hypo-, and stable methylation patterns with age were observed in males at these sites (**Figure 2**).

**Figure 2:**
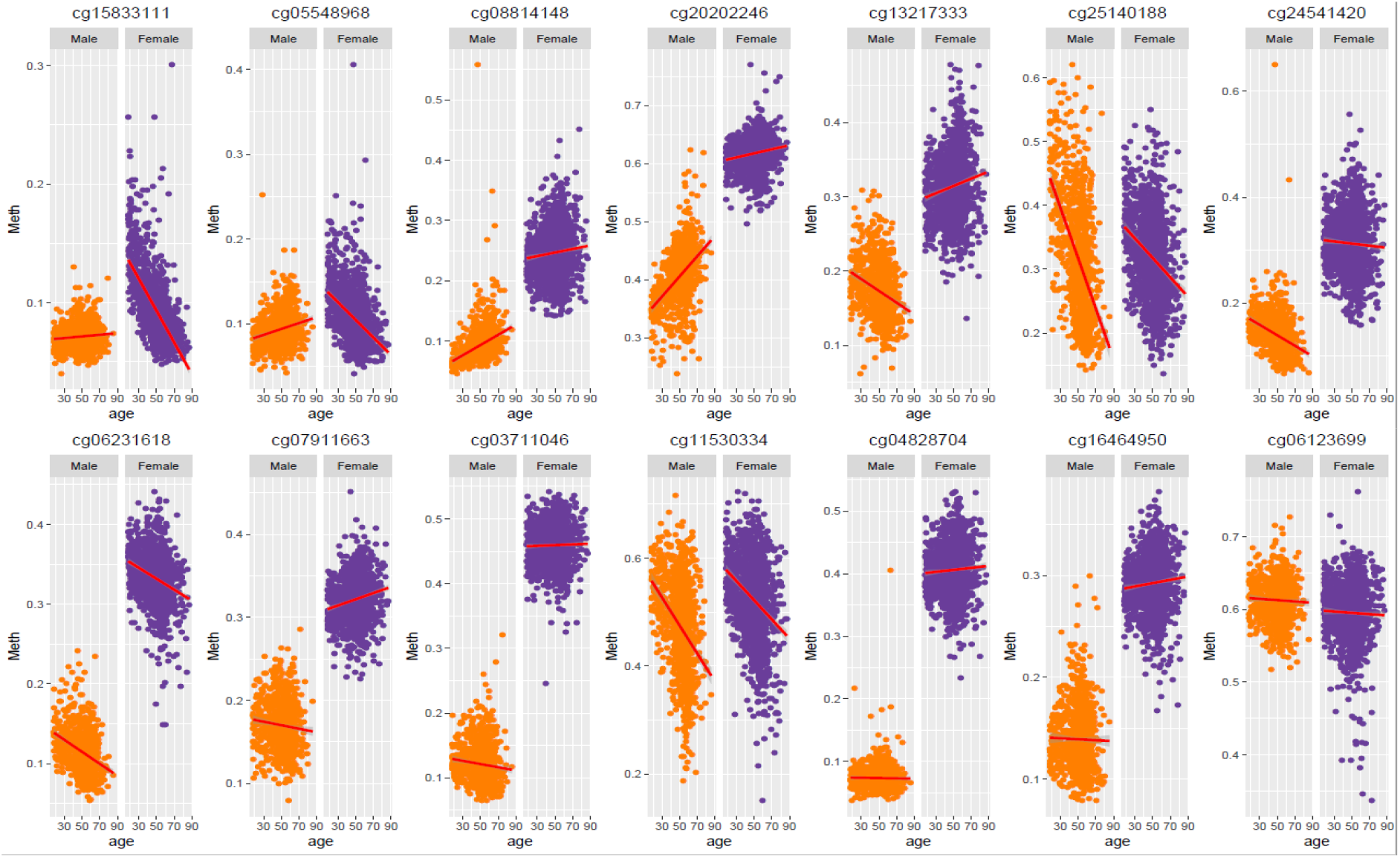
Correlation between DNA methylation and age in males versus females at 14 sites associated with the age-by-sex interaction

To investigate whether X-chromosome inactivation (XCI) affected the seven genes mapping to the 12 X-linked CpGs identified in this analysis, the genes were queried against a list of 114 XCI-escaping genes [19]. One gene, *RENBP*, was present in the list suggesting the majority of age-by-sex associated sites on the X chromosome are within genes that undergo XCI. The overlap between age-by-sex-associated genes and XCI escapees was not significant (χ^2^ = 0.07; P = 0.791). DNAm at the age-by-sex associated probes was generally weakly correlated with probes annotated to the same gene (**Supplementary Figures 2–8**), with the exception of *ANKRD58* (maximum correlation = 0.75) and *TMEM164* (maximum correlation = 0.83), in which correlations were present with nearby probes within 1500 bp of the gene transcription start site.

The seven age-by-sex-associated genes were queried against published GWAS results to determine whether there was enrichment among specific traits. One gene, *RENBP*, harboured a SNP variant (rs2269372, chr X:153,207,545; 2kb from the sexually-dimorphic CpG site) that was previously associated with schizophrenia in a Han Chinese population, and systemic lupus erythematosus in a European population, at a genome-wide significance threshold of P < 5×10^−8^ [20, 21, 22]. Of the 14 probes significantly associated with the age-sex interaction, eight were also present on the Illumina 450k array (all of which were X-linked), and were therefore queried against the ARIES mQTL database (mQTLdb; [23]). Three *trans*-associations were present among middle-aged individuals between cg04828704 and intronic variants in the chromosome 17 gene *PITPNC1* (rs2642065, rs2642041 and rs2642039; **Supplementary Table 2**). Two *trans*-associations were present between an intergenic X-linked probe (cg08814148), and two intergenic variants on chromosome 13 (rs113706784 and rs61957632)

### Gene expression analysis

To determine whether the observed sex differences in age-associated DNAm are recapitulated at the RNA level, the relationship between age intervals and gene expression (of the seven significant genes highlighted in the age-by-sex interaction model) was assessed in whole blood-derived samples from GTEx [17]. Spearman correlations were assessed between age and gene expression separately in males and females, with linear regression models used to formally test for significant age-by-sex interactions, using the median age from each binned age group. With the exception of *GAGE10*, all age-by-sex-associated genes were present in the blood-based gene expression dataset. There was no evidence for a significant age-by-sex interaction for expression levels in *CXorf38, RENB, OTUD5 or MID2* (P-value range 0.25 – 0.69). Significant age-by-sex interactions were present in the expression of *ANKRD58* (Beta = −0.02, P = 0.03) and *TMEM164* (Beta = −0.02, P = 0.02). However, these associations were attenuated following correction for multiple tests. The gene with the greatest absolute difference in age-by-expression correlations between males and females was *TMEM164* (r_male_ = −0.19, P_male_= 0.003; r_female_ = −0.40, P_female_= 3 × 10^−6^). There was a significant positive correlation between age and expression of *OTUD5* in males, but not in females (r_male_ = 0.26; P_male_ = 6.0 × 10^−5^; r_female_ = 0.17; P_female_ = 0.08). *ANKRD58* expression was significantly negatively correlated with chronological age in both males and females (r_male_ = −0.14; P_male_ = 0.03; r_female_ = −0.34; P_female_ = 9.0 × 10^−5^), whereas *MID2* was significantly positively correlated with chronological age in both sexes (r_male_ = 0.21; P_male_ = 9.7×10^−4^; r_female_ = 0.25; P_female_ = 0.005). As *GAGE10* expression data were not available in the whole blood-derived samples, age-associated expression trajectories were investigated in ovary and testis samples. There was a significant positive relationship between *GAGE10* expression and chronological age in ovarian samples (r = 0.25; P = 0.006; **Figure 3**), whereas expression remained stable with age in males (r = 0.05; P = 0.44; **Figure 3**). However, these correlations did not differ significantly from one another (P =0.08).

**Figure 3.**
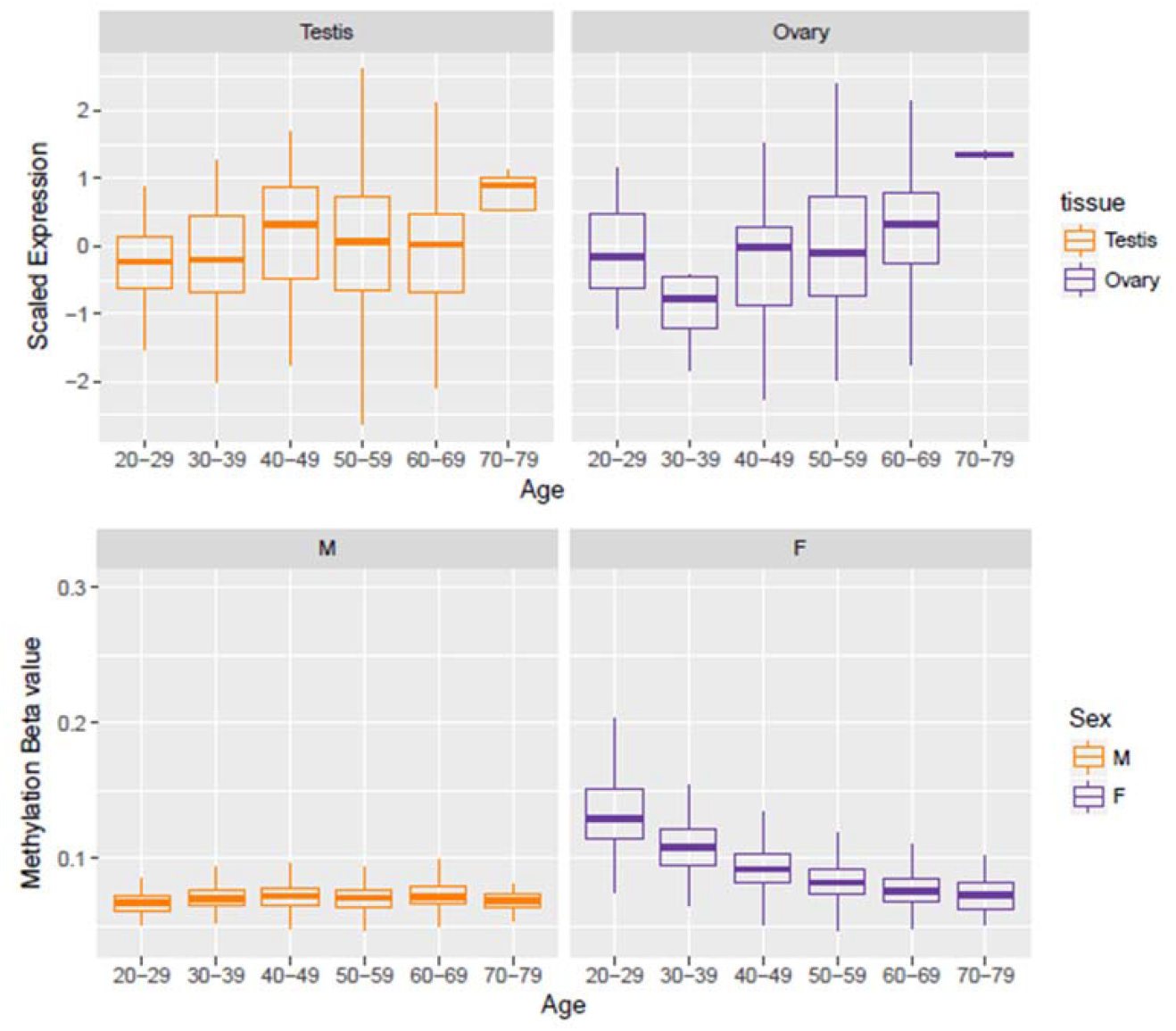
*GAGE10* gene expression and DNA methylation vs chronological age in males and females.

### Functional enrichment analysis

The genes where differential methylation was associated with chronological age and the age-by-sex interaction were assessed for enrichment in KEGG pathways, GO terms and tissue-specific expression in 30 general tissues from GTEx v7 [17, 24].

Two KEGG pathways, “cAMP signaling pathway” and “Transcriptional misregulation in cancer” were enriched for age-associated differentially-methylated genes (Bonferroni-adjusted P < 0.05) whereas 48 GO terms were enriched for these genes after correction for multiple tests (**Supplementary Table 3**). The age-associated genes were significantly enriched among differentially-expressed genes in 28 of 30 general tissues from GTEx, with no significant enrichment among differentially-expressed genes in fallopian tube and testis (**Supplementary Figure 9**).

There were no KEGG pathways enriched for the genes associated with the age-by-sex interaction. No GO terms were enriched for these genes after correction for multiple testing (Bonferroni-adjusted P < 0.05). However, four terms relating to protein dimerization and binding were nominally significantly associated with these genes (P < 0.05; **Supplementary Table 4**). There was significant enrichment for age-by-sex-associated genes among downregulated differentially-expressed genes in the lung, liver, testis and blood (**Figure 4**).

**Figure 4:**
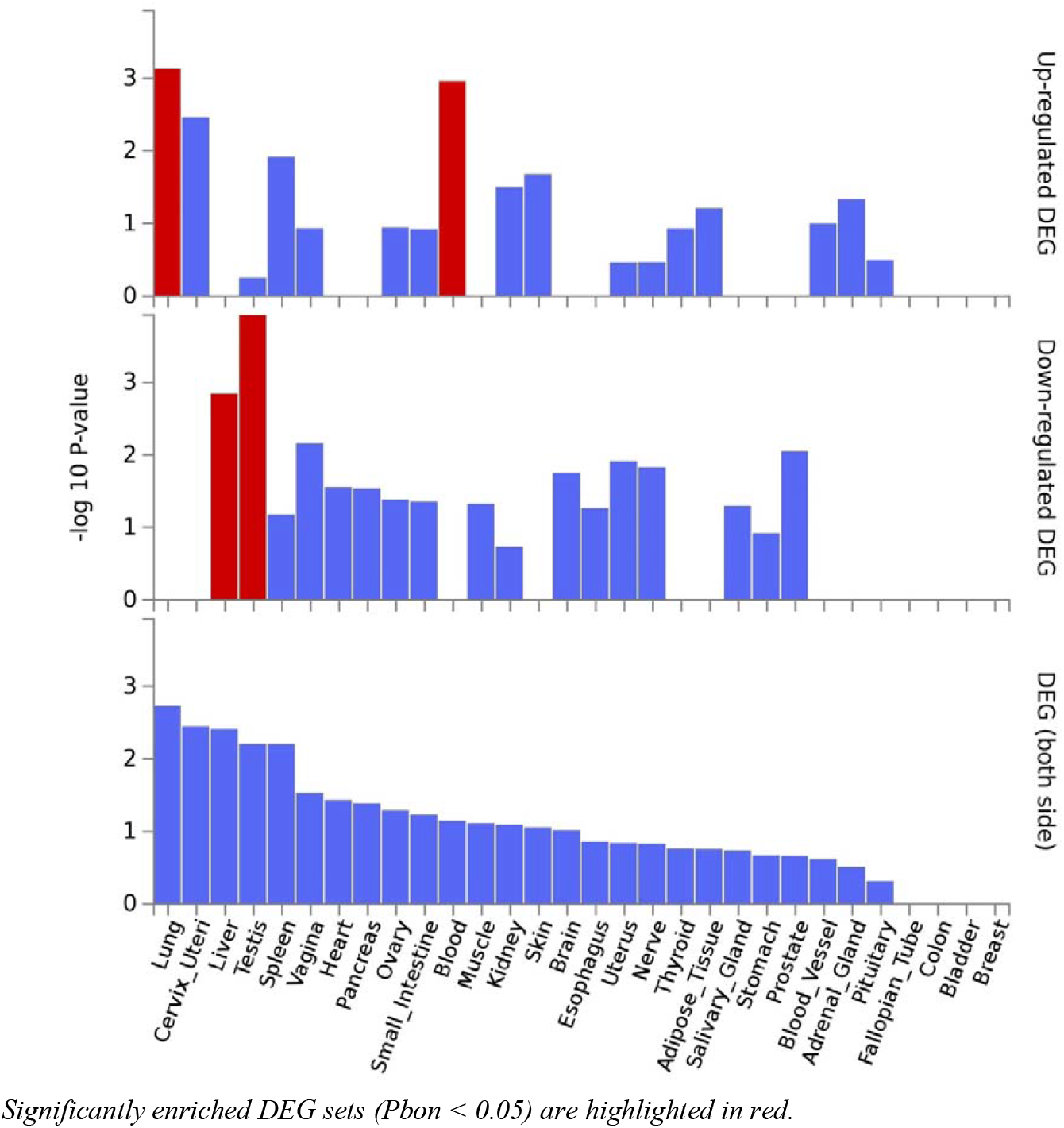
Enrichment of age-by-sex associated genes among differentially-expressed genes in 30 general tissue types (GTEx)

## Discussion

We identified 14 loci displaying sexually-dimorphic ageing associations in DNAm across the adult age range. These were predominantly located on the X-chromosome, with the exception of two loci mapping to chromosomes 3 and 8, and mapped to seven genes, all on the X chromosome. The DNAm correlation structure across these genes suggests that the age-associated methylation trajectories at the sites identified in this study do not reflect a gene-wide modification of XCI, but rather localised changes. Where strong correlations with age-by-sex-associated probes were observed, these were within 1500 bp of the TSS.

The site with the greatest absolute age-DNAm correlation difference between males and females mapped to *GAGE10*, a member of the *GAGE* cancer/testis antigen family [25].

Normal expression of GAGE proteins is limited to germ cells. However, *GAGE* transcripts have been observed in multiple cancers including melanomas, breast, lung, ovarian and thyroid cancers [26, 27, 28, 29, 30, 31].

Six of the seven age-by-sex-associated genes did not return results when queried in the GWAS catalogue. This is not surprising as the X chromosome is usually omitted from association studies [32]. However, one of the age-by-sex-associated genes, *RENBP5*, has been associated with traits by GWAS (schizophrenia [Han Chinese population] and systemic lupus erythematosus [European ancestry]) [21, 22]. The incidence rates of both disorders differ between males and females. Schizophrenia is more common in males, who are also more likely to experience negative symptoms than females (e.g. anhedonia) [33], whereas systemic lupus erythematosus has a female to male ratio of disease incidence of 9:1 [34].

Analysis of tissue-specific expression patterns of the seven genes displaying sexually-dimorphic age-associated DNAm trajectories revealed an enrichment among differentially-expressed genes in liver, testis, lung and blood. Enrichment among differentially-expressed genes in testis and liver may be indicative of a relationship with endocrine function and sex-specific ageing. This is consistent with current hypotheses of endocrine differences as a contributor to the disparity in the life expectancy of males and females [35, 36, 37].

Three *trans*-SNP-CpG associations were present among the 14 sites associated with the age-by-sex interaction, and were within an intronic region of the chromosome 17 gene *PITPNC1*. These were associated with DNAm at an X-linked locus mapping to *MID2*. Differential expression of *MID2* has been reported in breast cancer, with increased levels associated with poorer prognostic outcomes [38]. Similarly, overexpression of *PITPNC1* has been found to promote metastasis in multiple types of cancer [39].

The mQTLdb resource is limited to probes present on the Infinium Human Methylation 450K BeadChip: the predecessor of the Infinium MethylationEPIC BeadChip used in the current study [23]. Of the 14 sites queried for mQTLs, eight were present on both platforms. It is, therefore, possible that additional, as-yet unidentified SNP associations are present among the remaining 12 sites specific to the Infinium MethylationEPIC BeadChip.

In addition to the age-by-sex interaction, we examined the relationship between DNAm and chronological age. We replicated previous findings, with the strongest age-associated effects observed in *KLF14, ELOVL2* and *FHL2* [4, 5, 18].

A limitation of this study is the absence of gene expression data for the Generation Scotland cohort. Publicly-available gene expression data were used to compare age-associated expression and DNAm patterns; however, it is not possible to determine how comparable the age distributions are between the Generation Scotland DNAm dataset and the GTEx gene expression dataset, where sample ages are presented in intervals. The current study is strengthened by the use of a large, unrelated cohort with a broad age range, which has permitted the development of pseudo-longitudinal profiles of DNAm across the life course in males and females.

In conclusion, we have identified 14 CpG sites displaying differences in age-associated DNAm patterns between males and females. Twelve of these sites are located on the X-chromosome, two within genes which have been linked to multiple cancers, and one in a gene which has been implicated in both schizophrenia and systemic lupus erythematosus by GWAS. In order to identify mechanisms of sex-specific differences in biological ageing, further investigation of sexual dimorphic characteristics of these processes over the life course is warranted.

---

**Supplementary Figure 1:**
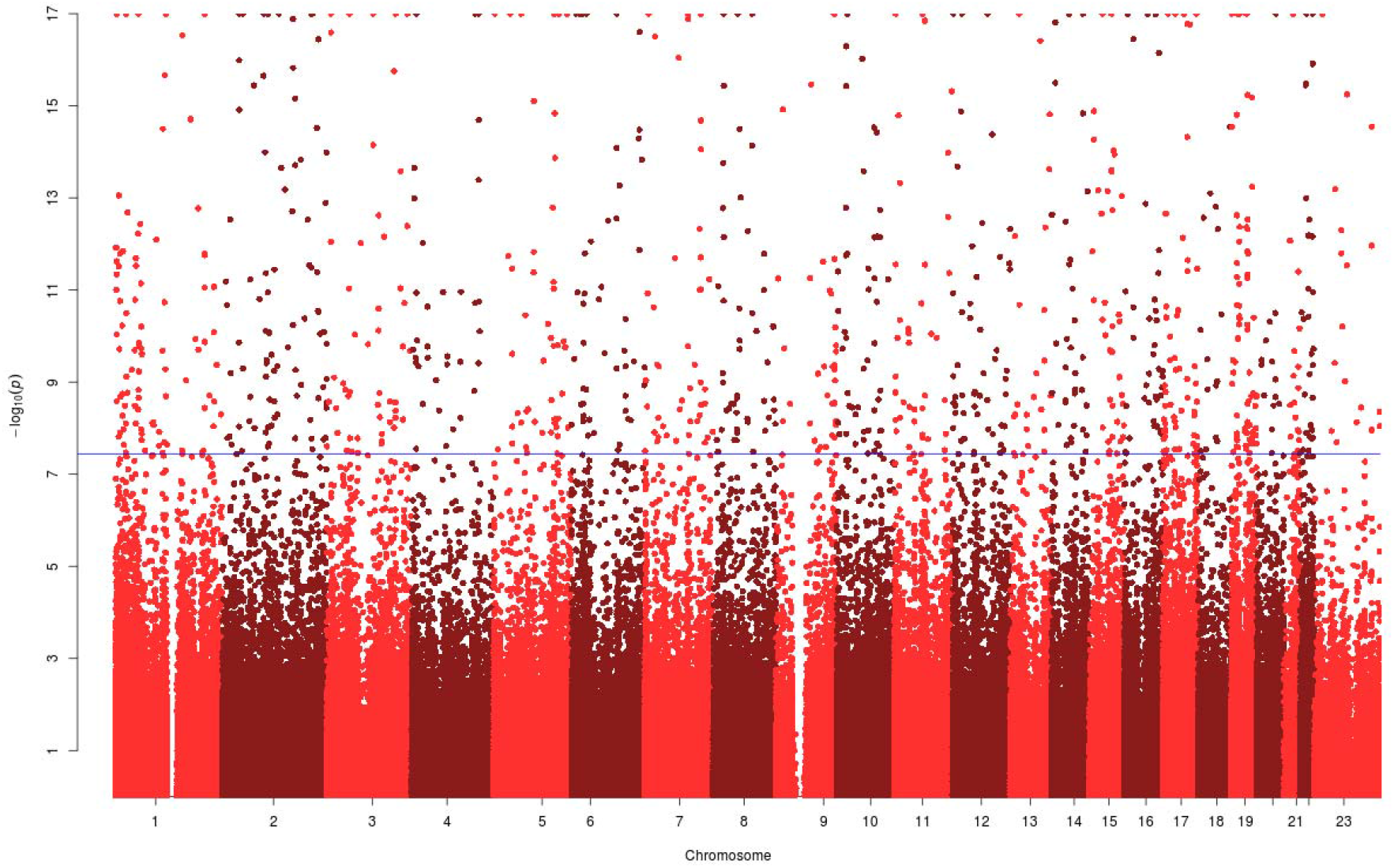
Manhattan Plot for the EWAS of chronological age in Generation Scotland. The −log_10_(p value) has been truncated at 17.

**Supplementary Figure 2:**
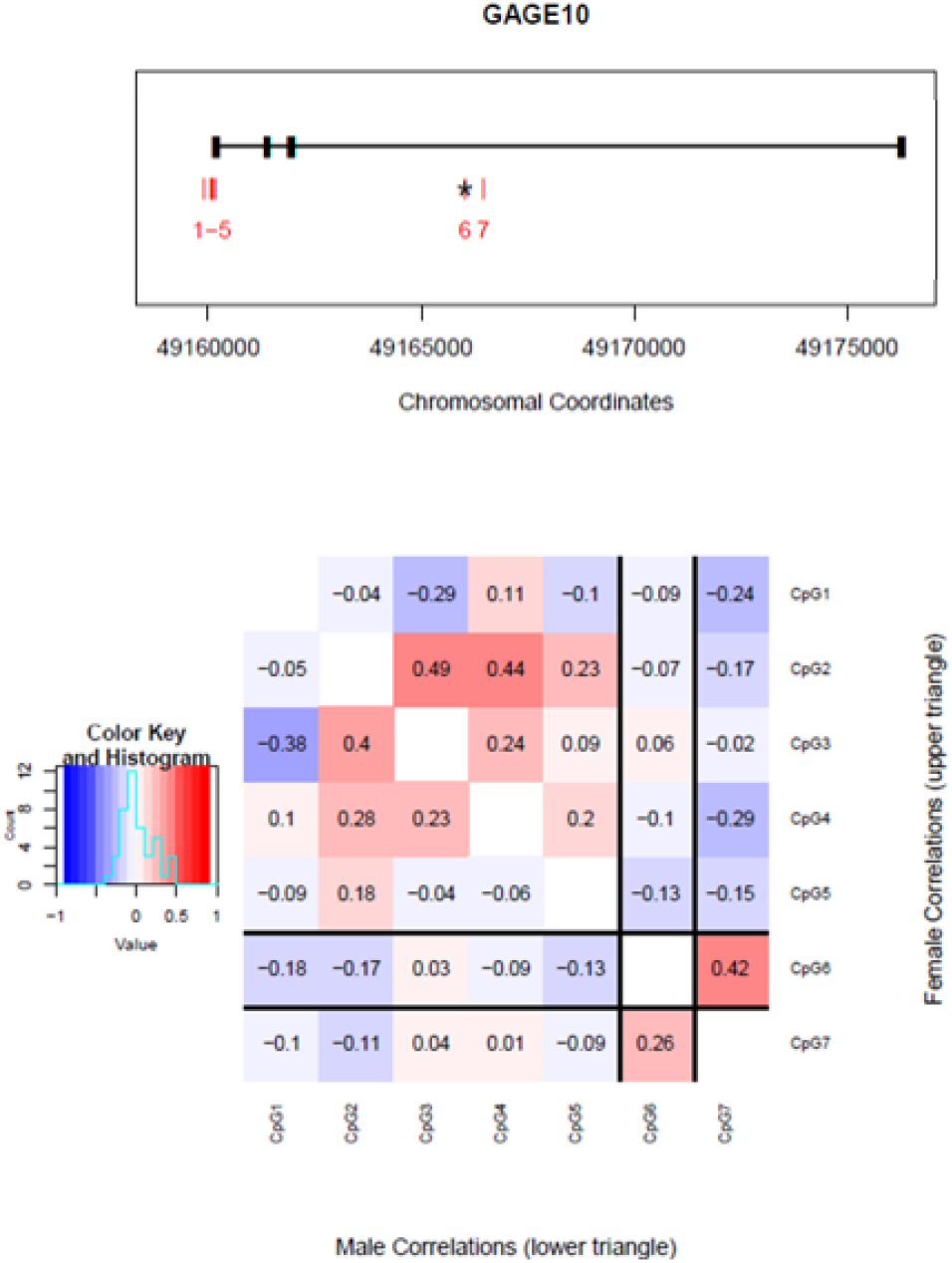
*GAGE10* DNA methylation summary

**Supplementary Figure 3:**
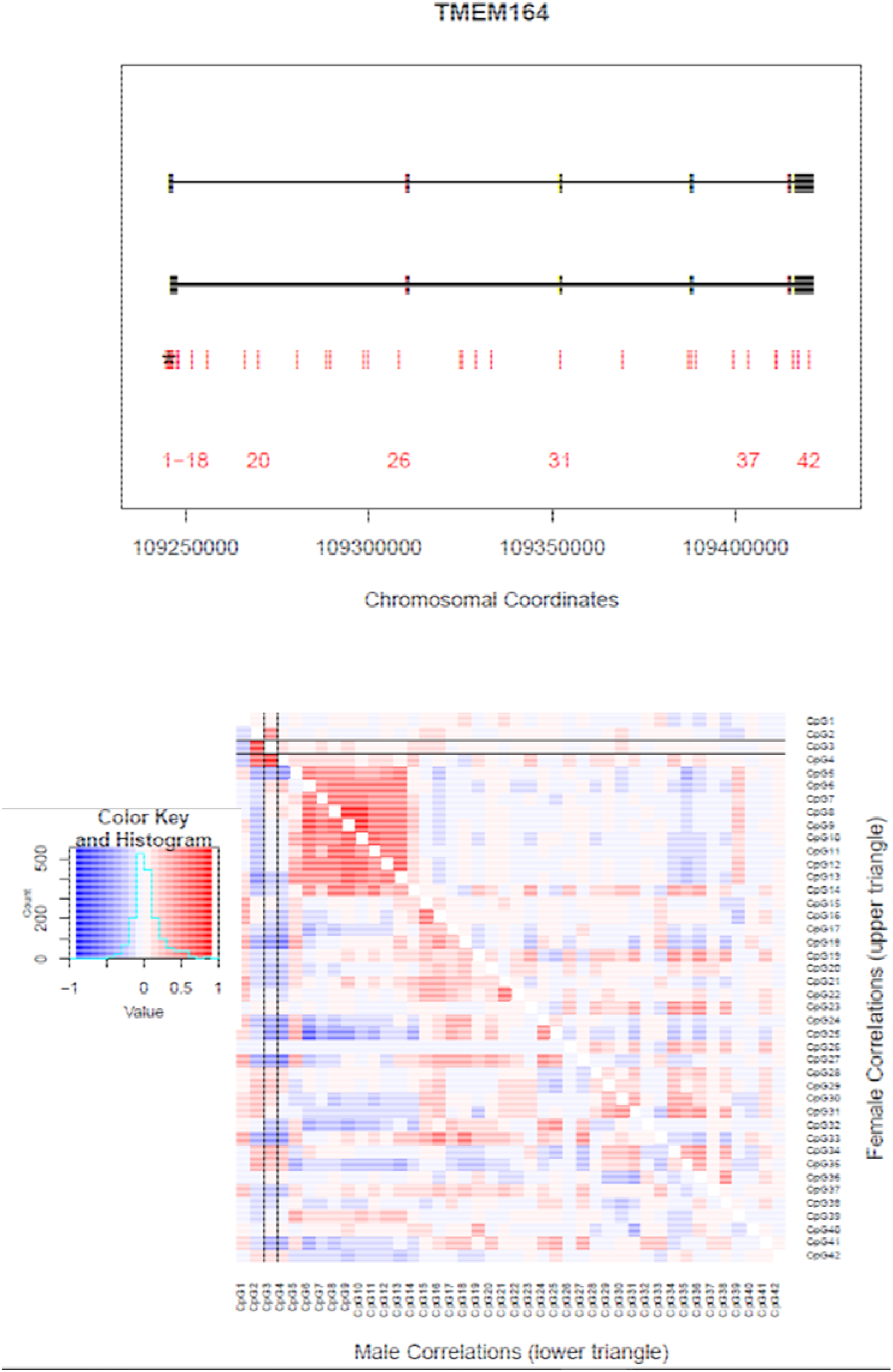
*TMEM164* DNA methylation summary

**Supplementary Figure 4:**
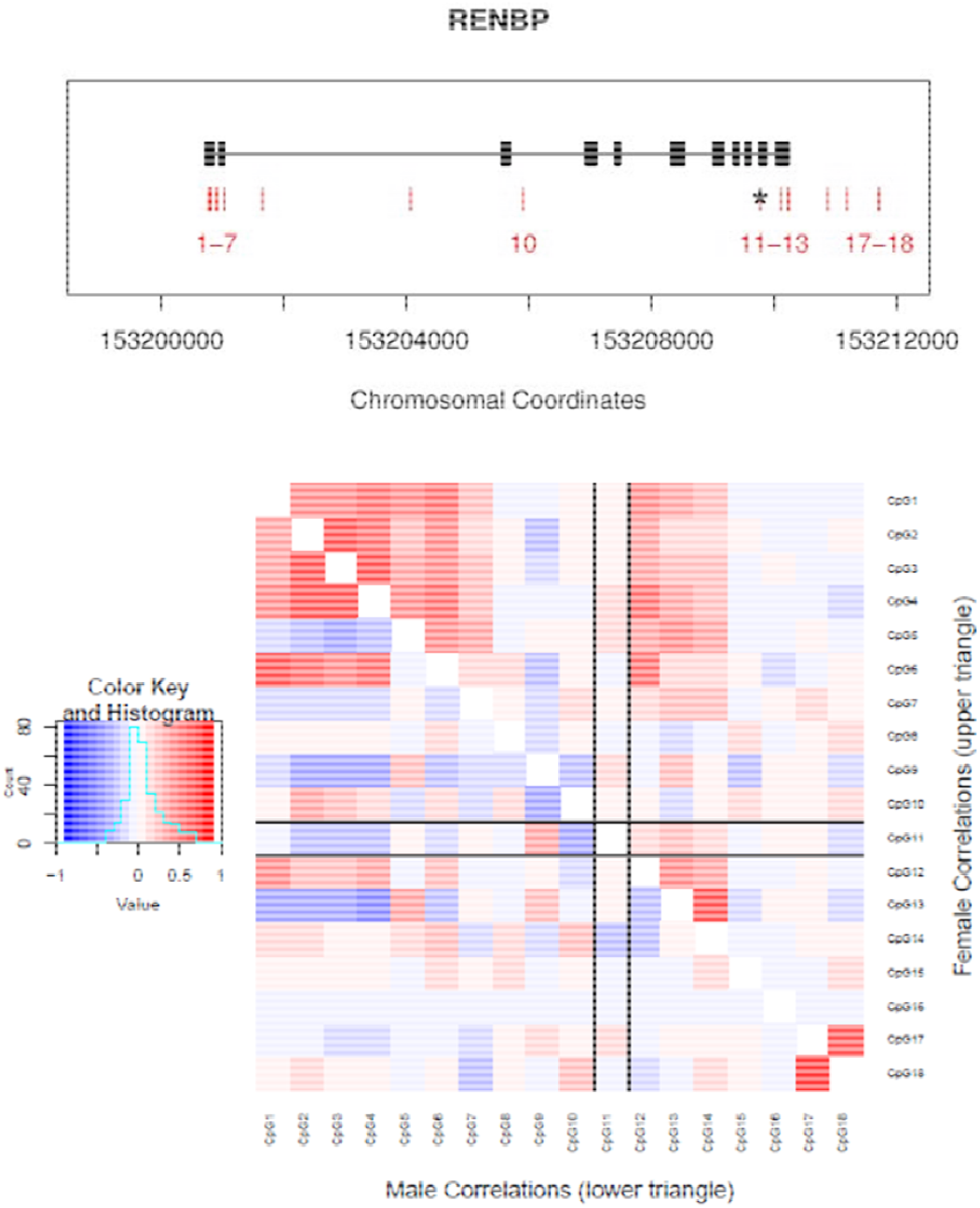
*RENBP* DNA methylation summary

**Supplementary Figure 5:**
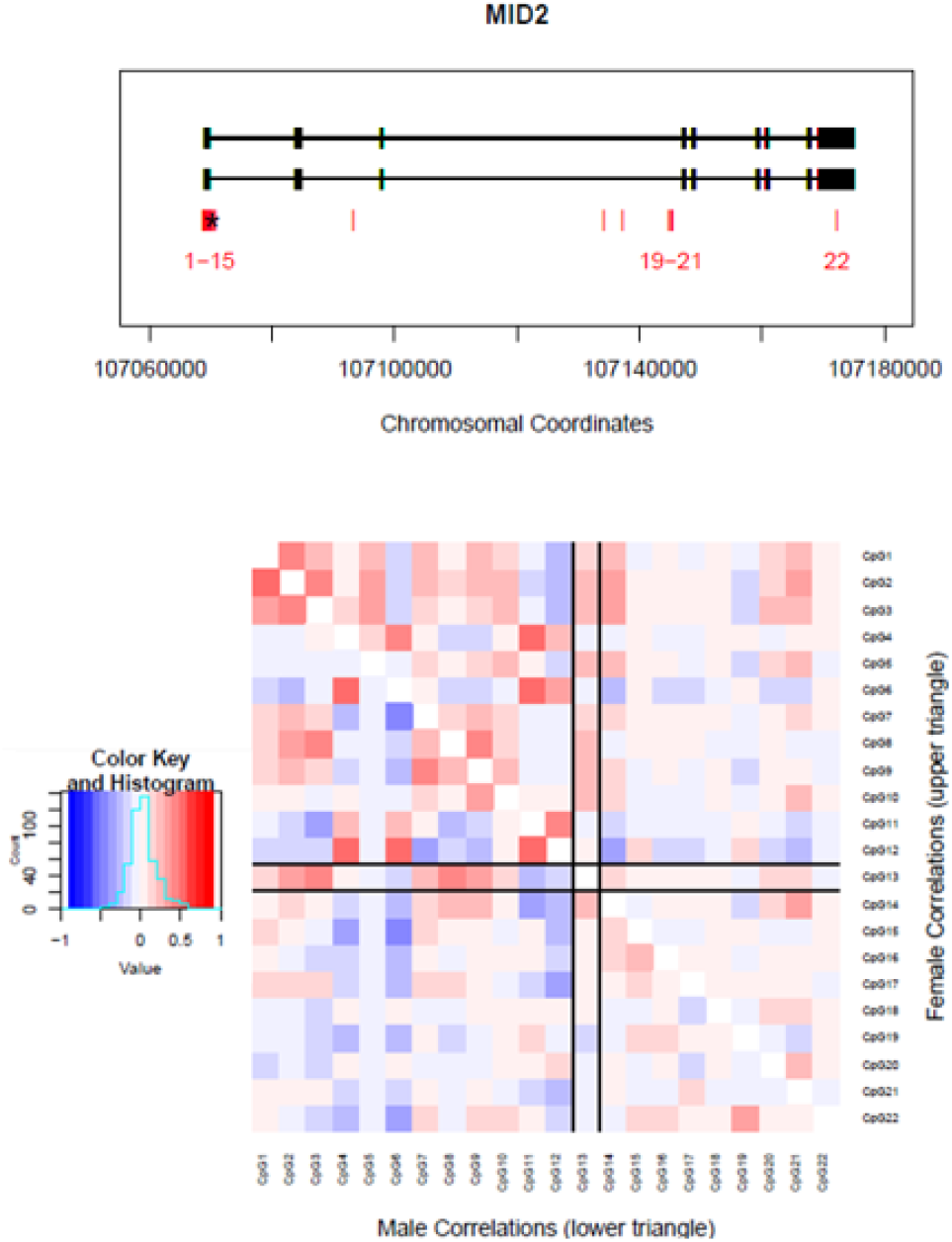
*MID2* DNA methylation summary

**Supplementary Figure 6:**
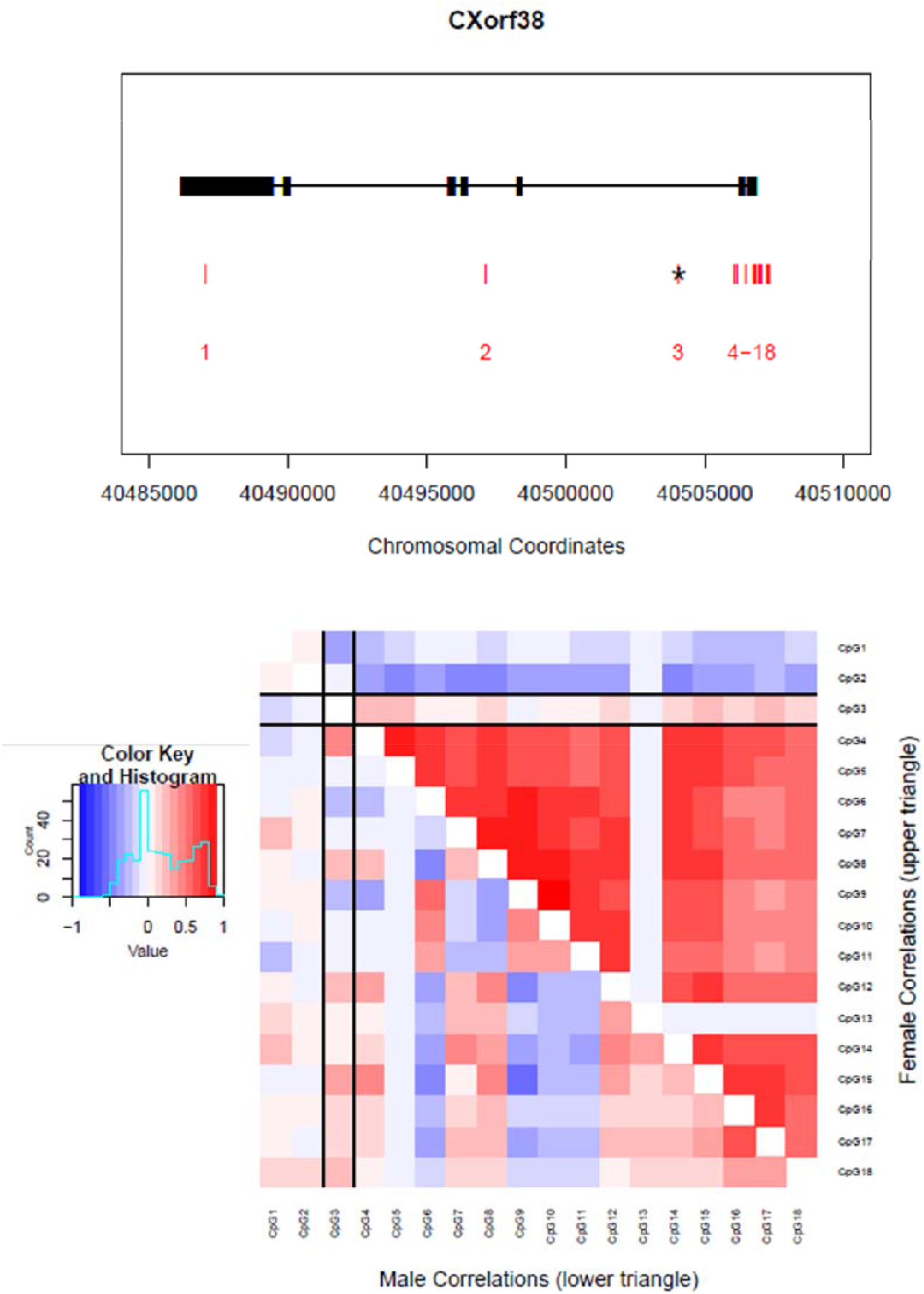
*CXorf38* DNA methylation summary

**Supplementary Figure 7:**
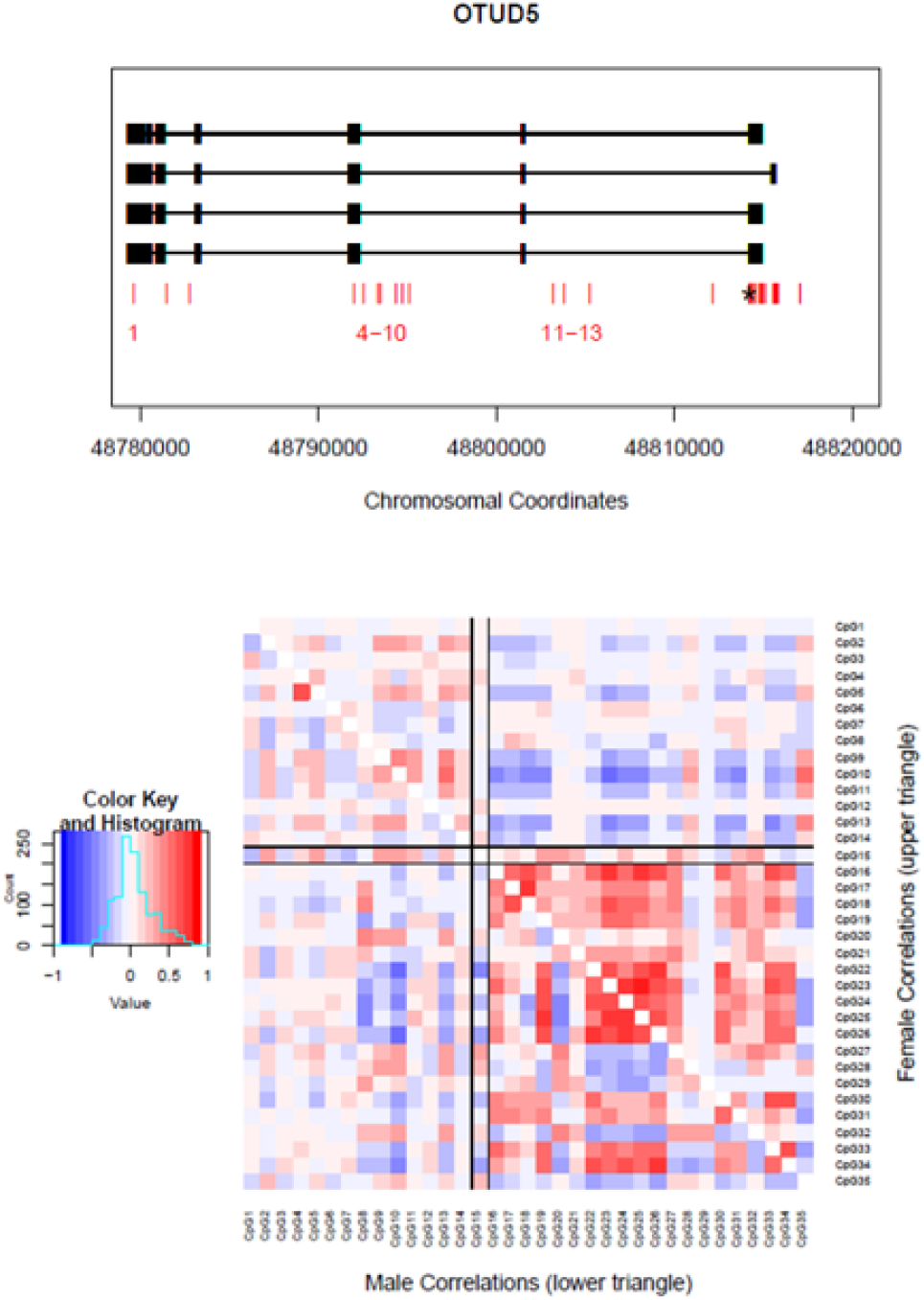
*OTUD5* DNA methylation summary

**Supplementary Figure 8:**
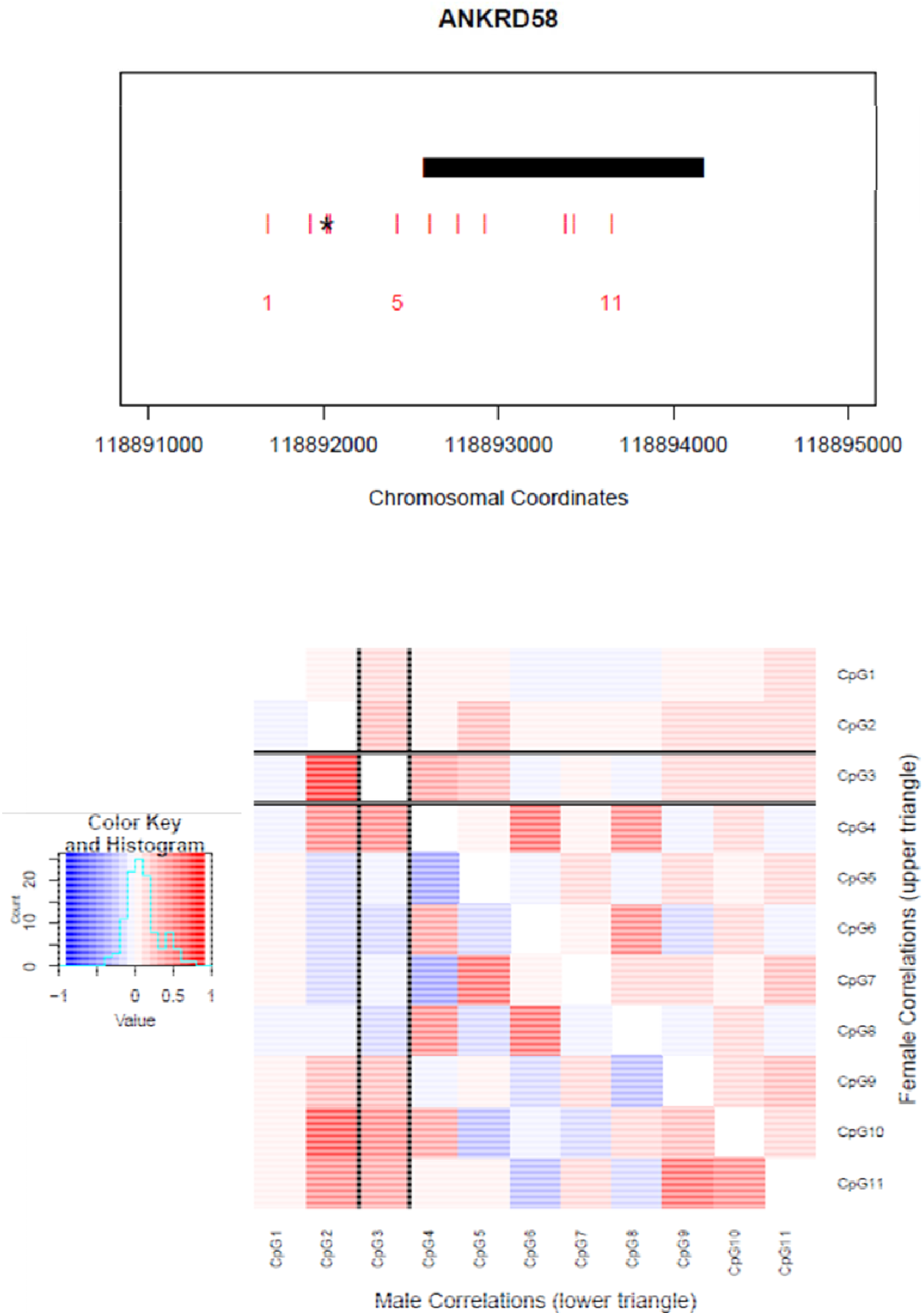
*ANKRD58* DNA methylation summary

**Supplementary Figure 9:**
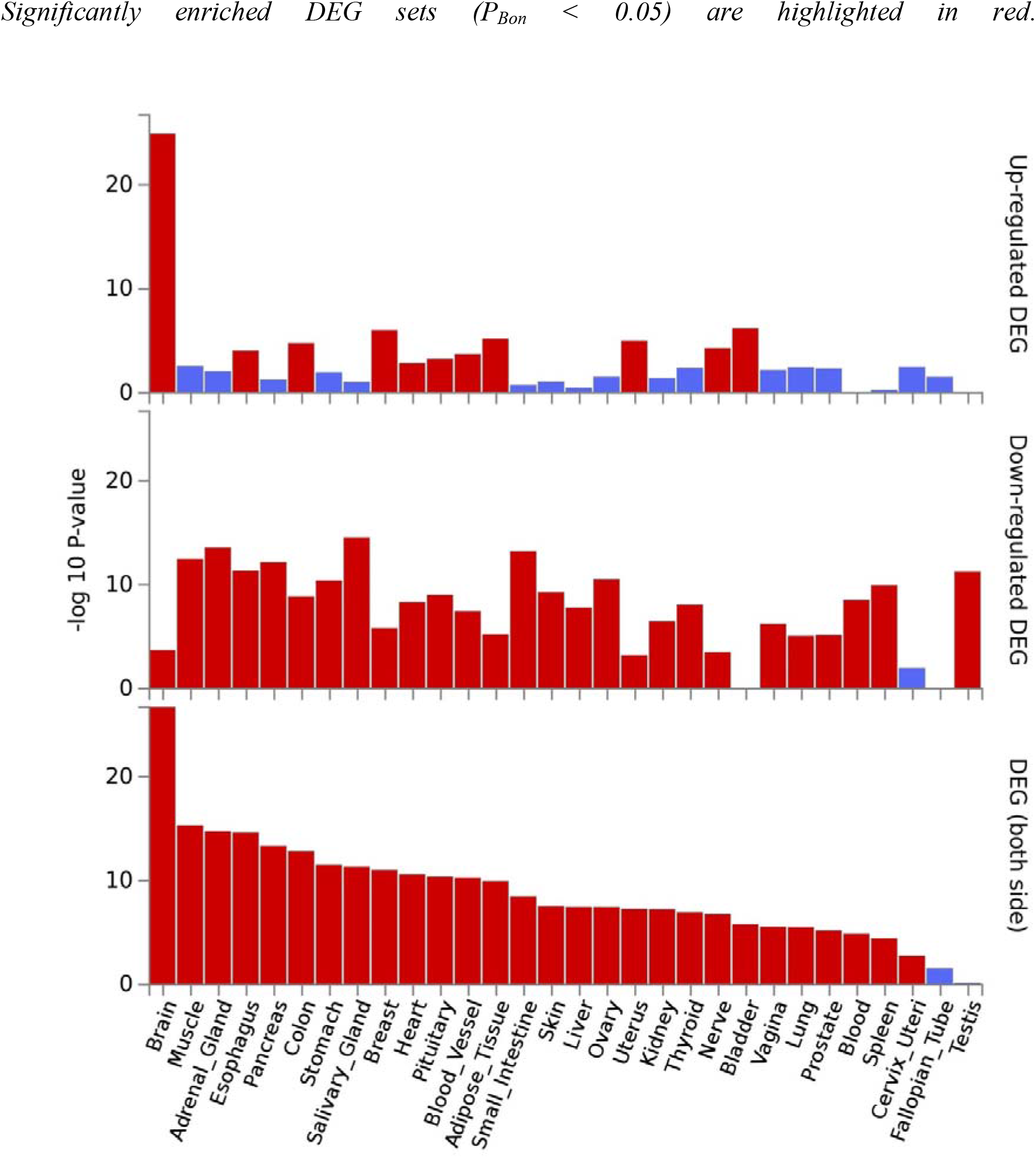
Enrichment of age associated genes among differentially-expressed genes in 30 general tissue types (GTEx)

**Supplementary Table 2:** Middle age SNP-CpG associations reported for probes associated with the age-by-sex interaction

| SNP | SNP Chr | SNP Pos | SNP Gene | A1 | A2 | MAF | CpG | CpG Chr | CpG Pos | CpG Gene | beta | t-stat | Effect Size | P |
| --- | --- | --- | --- | --- | --- | --- | --- | --- | --- | --- | --- | --- | --- | --- |
| rs113706784 | 13 | 59106653 | Intergenic | C | T | 0.425 | cg08814148 | X | 118407645 | Intergenic | 0.246 | 5.396 | 0.005 | 0.246 |
| rs61957632 | 13 | 59106880 | Intergenic | T | C | 0.411 | cg08814148 | X | 118407645 | Intergenic | 0.264 | 5.735 | 0.006 | 0.264 |
| rs2642065 | 17 | 65638272 | *PITPNC1* | T | G | 0.209 | cg04828704 | X | 107070200 | *MID2* | -0.281 | -5.613 | 0.008 | -0.281 |
| rs2642039 | 17 | 65644419 | *PITPNC1* | G | A | 0.243 | cg04828704 | X | 107070200 | *MID2* | -0.252 | -5.384 | 0.004 | -0.252 |
| rs2642041 | 17 | 65645744 | *PITPNC1* | T | C | 0.212 | cg04828704 | X | 107070200 | *MID2* | -0.266 | -5.400 | 0.001 | -0.266 |

**Supplementary Table 3:**
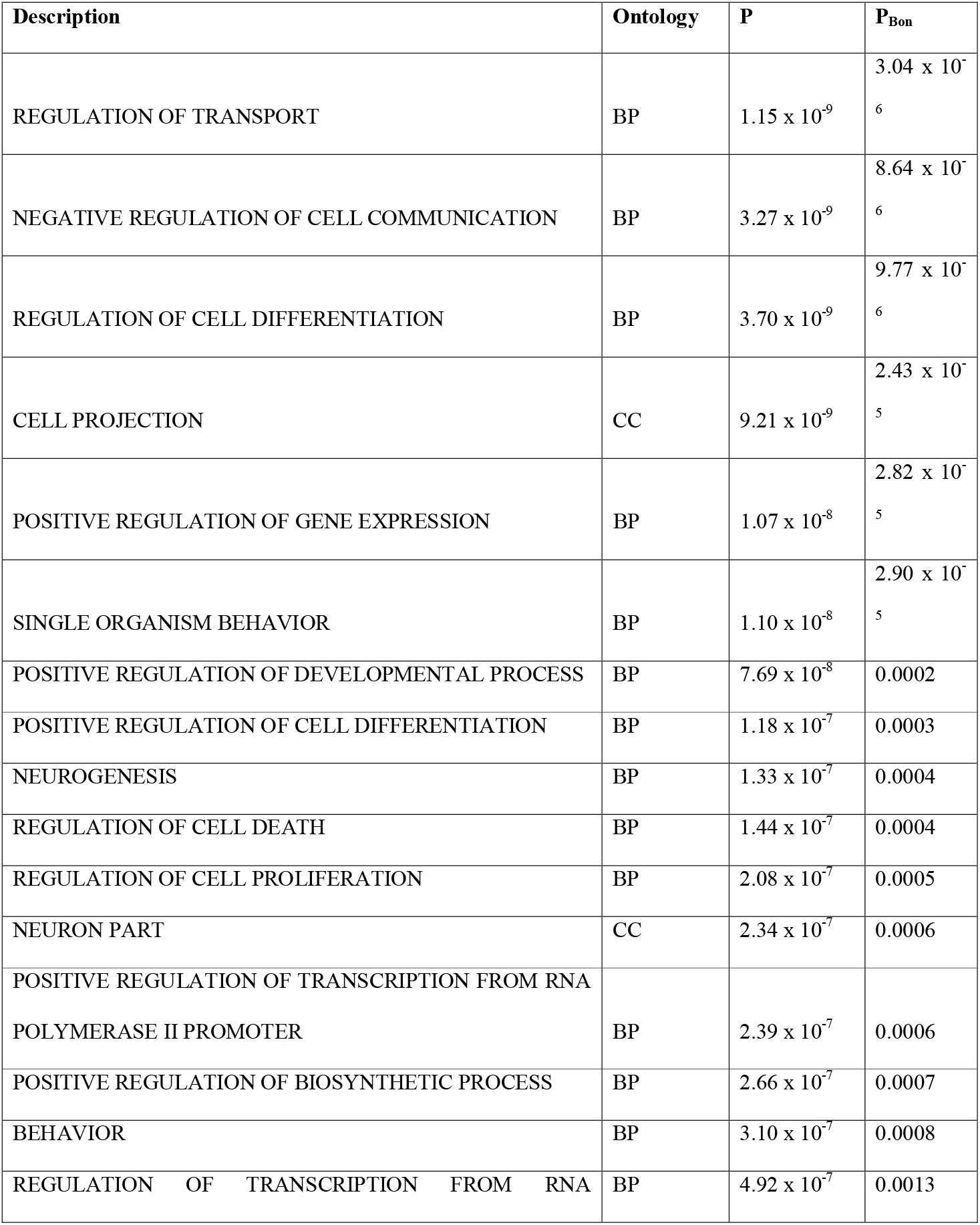
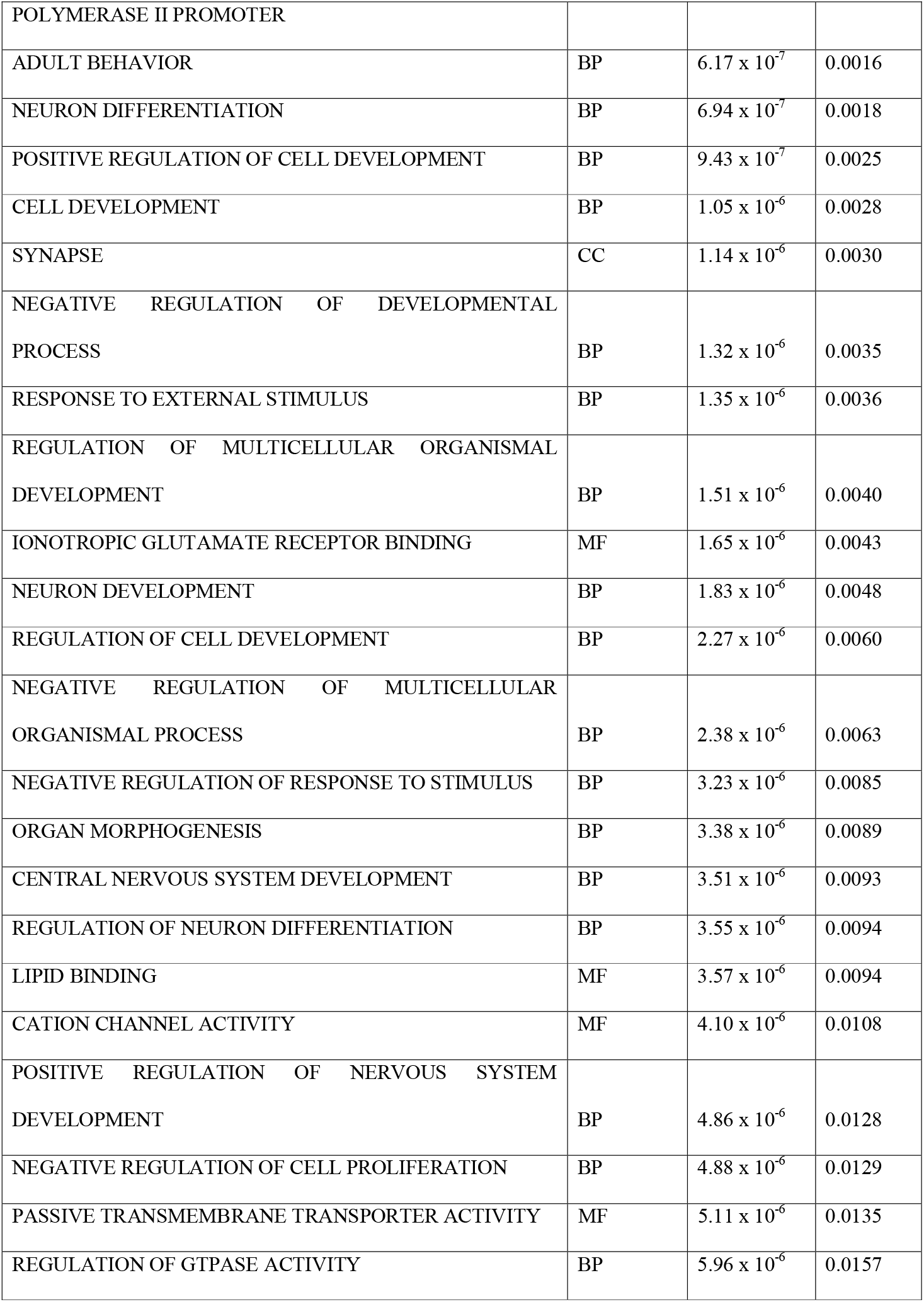
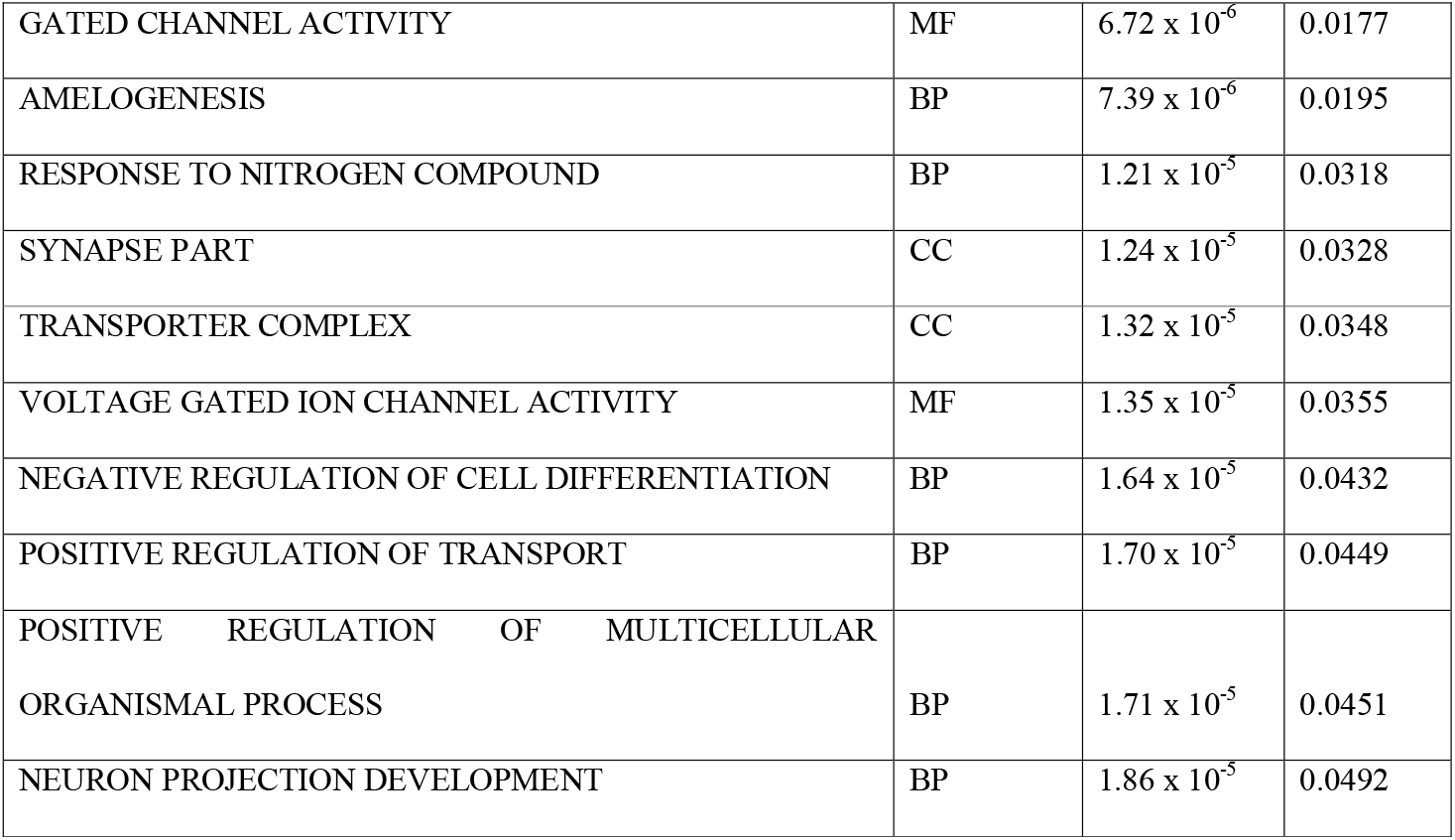
GO Terms enriched for genes showing age associated differential methylation

**Supplementary Table 4:** GO Terms enriched for genes showing age-by-sex associated differential methylation

| Description | Ontology | P | P<sub>Bon</sub> |
| --- | --- | --- | --- |
| PROTEIN HOMODIMERIZATION ACTIVITY | MF | 0.02 | 0.06 |
| PROTEIN MODIFICATION BY SMALL<br>PROTEIN CONJUGATION OR REMOVAL | BP | 0.02 | 0.09 |
| PROTEIN DIMERIZATION ACTIVITY | MF | 0.04 | 0.15 |
| IDENTICAL PROTEIN BINDING | MF | 0.04 | 0.16 |

